# Rosiglitazone in the thawing medium improves mitochondrial function in stallion spermatozoa through Akt phosphorylation and reduction of caspase 3

**DOI:** 10.1101/532689

**Authors:** JM Ortiz-Rodriguez, C Balao da Silva, J Masot, E Redondo, A Gazquez, MC Gil, C Ortega Ferrusola, FJ Peña

## Abstract

**Background:** The population of stallion spermatozoa surviving thawing, experience, among other changes, compromised mitochondrial functionality and accelerated senescence. It is known that the stallion spermatozoa show very active oxidative phosphorylation that may accelerate sperm senescence through increased production of reactive oxygen species. Rosiglitazone has proven to enhance the glycolytic capability of stallion spermatozoa maintained in refrigeration.

**Objectives:** Thus, we hypothesized that thawed sperm may also benefit from rosiglitazone supplementation.

**Material and Methods:** Thawed sperm doses were washed and re-suspended in Tyrodes media, split sampled and supplemented with 0 or 75 μM rosiglitazone. After 1 and two hours of incubation, mitochondrial functionality, Akt phosphorylation and caspase 3 activity were evaluated. Further samples were incubated in presence of the Akt1/2 inhibitor, compound C (AMPK inhibitor) and GW9662 (antagonist of the PPARγ receptor).

**Results:** Rosiglitazone maintained Akt phosphorylated and reduced caspase 3 activation (p<0.01) that was prevented by incubation in presence of the three inhibitors. Rosiglitazone also enhanced mitochondrial functionality (p<0.01).

**Conclusion:** We provide, for the first time, evidences that the functionality of frozen stallion spermatozoa can be potentially improved after thawing through the activation pro survival pathways, opening new clues to improve current sperm biotechnologies.

## INTRODUCTION

The stallion spermatozoa can be stored in the liquid state for short periods, or frozen for long-term storage. Frozen thawed sperm has numerous advantages, however its wider use is still constrained by a number of weaknesses [1]. Among them, the high stallion-to-stallion variability and the insufficient standardization of the freezing and thawing procedures. While cryopreservation induces mortality to on average 50% of the initial sperm population, the surviving spermatozoa are not completely functional; on the contrary they experience accelerated senescence that requires more intense and costly mare management for insemination to compensate this reduced lifespan. Most of the studies on sperm cryopreservation have aimed to increase the number of spermatozoa surviving the procedure, but studies aiming to improve the quality of the surviving population are scarce. Although the changes induced by cryopreservation have been extensively investigated, mostly focusing on cryopreservation induced necrosis [2–4], few studies have addressed the physiology of spermatozoa surviving freezing and thawing. There are not many studies that have tried to develop measures to increase the quality of frozen sperm after thawing, with the exception of procedures to remove dead and damaged spermatozoa from the cryopreserved sample [5–7]. The population of spermatozoa surviving freezing and thawing, experience changes recently termed spermptosis [8]. These changes basically represent an acceleration of the default apoptotic dead that most of the spermatozoa experience after ejaculation [9, 10]. In brief, osmotic shock induces membrane and mitochondrial damage [11], the mitochondrial damage causes impairment of redox regulation, leading to lipid, protein and DNA modifications in the spermatozoa, resulting in decreased motility and viability [8, 12, 13]. Other changes recently described in relation to cryopreservation include increased intracellular Na+ and membrane depolarization due to the compromised functionality of the Na+-K+ATPase pump [14]. One key protein related to sperm survival is the Akt or protein kinase B. Provided that this kinase is phosphorylated, spermatozoa remain viable, but upon dephosphorylation of Akt, pro-caspase 3 is cleaved and the spermatozoa rapidly lose motility [15, 16]. The cryopreserved semen show impaired mitochondrial activity due to oxidative stress and the osmotic damage occurring at thawing [17–20]. This damage compromises the capability of thawed stallion spermatozoa to produce ATP through oxidative phosphorylation. Thus, the cryopreserved sperm has lower motility and lower ATP content compromising their functionality, and finally their fertilizing ability. Rosiglitazone has proven to enhance the glycolytic capability of stallion spermatozoa maintained in refrigeration for long periods [21], moreover human [22] and porcine [23] studies indicate that rosiglitazone activate pro-survival pathways in spermatozoa. In view of these facts, we hypothesized that thawed stallion spermatozoa may respond to rosiglitazone and thus rosiglitazone supplementation could be a suitable strategy to improve the quality of thawed stallion spermatozoa.

## MATERIAL AND METHODS

### Reagents and media

Hoechst 33342 (Excitation: 350 nm, Emission: 461 nm) (Ref: H3570); 5,5’,6,6’-tetrachloro-1,1’,3,3’tetraethylbenzymidazolyl carbocianyne iodine (JC-1), (Excitation: 488 nm; Emission: 530 nm, monomer form) (Excitation: 561 nm; Emission: 591 nm, aggregate form) (Ref: T3168); CellRox Deep Red Reagent (Excitation: 644 nm; Emission: 655nm) (Ref: C10422); Cell Event Caspase-3/7 Green Detection Reagent (Excitation, 502 nm; Emission: 530 nm) (Ref: C10423); Annexin V Alexa Fluor 647™ conjugate (Excitation: 650 nm, Emission: 665 nm) (Ref: A23204); ethidium homodimer (Excitation, 528 nm; Emission, 617 nm) (Ref: E1169); were purchased from ThermoFisher Scientific (Molecular Probes) (Waltham, Massachusetts, USA). Phospho-Akt (Ser 473) (D9E) XP^®^ Rabbit mAb (Alexa Fluor^®^ 488 conjugate was acquired from Cell Signalling technology (Danvers, Massachusetts, USA). Rosiglitazone, dorsomorphin, GW9662 and Akt I-II kinase inhibitor were purchased from Sigma-Aldrich (Madrid, Spain)

### Experimental design

Frozen doses of semen (6 different stallions) stored in our center were used in this study. Straws were thawed in a water bath at 37°C for at least 30 sec and diluted in pre-warmed INRA-96 extender to a final concentration of 50 x 10^6^ spermatozoa/ml. The samples were centrifuged (600 g x10’) and re-suspended in Tyrode’s media [24] to a final concentration of 50x10^6^ spermatozoa/ml. The semen was split in subsamples for control and experimental treatments and incubated in a water bath at 38°C. Initially a dose response experiment with rosiglitazone was performed and the minimally effective dose was chosen for subsequent experiments. Further experiments were conducted in presence of effective doses of rosiglitazone in presence of specific inhibitors. Sperm functions studied included motility and kinematics, mitochondrial membrane potential, production of superoxide and live spermatozoa, caspase 3, phosphorylation of Akt and determination of the oxidation reduction potential.

### Sperm motility

Sperm motility was assessed using a Computer Assisted Sperm Analysis (CASA) system (ISAS Proiser, Valencia, Spain). Semen was loaded in a Leja^®^ chamber with 20 μm of depth (Leja, Amsterdam, The Netherlands) and placed on a warmed stage at 37 °C. The analysis was based on an evaluation of 60 consecutive digitalized images obtained using a 10X negative phase-contrast objective (Olympus CX 41). At least three different fields were recorded to ensure that at least 300 spermatozoa were analyzed per sample. Spermatozoa with a VAP (average velocity) <15 μm/s were considered immotile, while spermatozoa with a VAP > 35 μm/s were considered motile. Spermatozoa deviating < 45% from a straight line were classified as linearly motile.

### Simultaneous detection of mitochondrial membrane potential and superoxide production

Semen aliquots were stained with JC-1 (90 nM) CellROX (2.5 μM) and HOECHST 33342 (0.25 μM) and incubated at r.t in the dark for 30 min [25]. The samples were then run in the flow cytometer and mitochondrial membrane potential and superoxide production were investigated only in live cells.

### Measurement of Oxidation-reduction potential

Oxidation-reduction potential was measured using RedoxSYS^®^ Diagnostic system (Englewood CO, USA). This is a novel technology that measures in 4 min the static oxidation reduction potential (sORP), measuring the potential of an electrochemical cell under static conditions, followed by measuring the capacity of oxidation reduction potential (cORP), which is the total amount of readily oxidizable molecules [26]. In brief, 30μL of semen was loaded in the sample port of the pre-inserted disposable sensor, the measurement beginning in this moment. After 4 minutes. The static oxidative-reduction potential (sORP) is provided in millivolts (mV). According to the manufacturer sORP are measured while applying a low oxidizing current (1nA) to the sample. After allowing 1 min and 50 s for equilibration the reader measures twice per second over 10 s the difference in potential between working and reference electrode in mV. Subsequently cORP are measured applying a linearly increasing oxidizing current until the charge rapidly changes between and working and reference electrode, indicating that all readily oxidizable molecules are oxidized. The time until the charge changes is used to calculate the number of electrons needed to cause charge changes and is reported in μCoulomb (μC).

### Staining for determination of live and dead cells and caspase 3 and 7 activity

This protocol has been developed in the laboratory where the present research was conducted and has been extensively described in previous publications [12, 13, 15, 27]. In brief, spermatozoa (5 x 10^6^/mL) in 1 mL of PBS were stained with CellEvent 2 μM and 0.5 μM Hoechst 33342 and incubated 25’ in the dark at r.t. Then 0.3 μM of Eth-1 was added to each sample and after incubation for 5 additional minutes, then samples were immediately evaluated in the flow cytometer.

### Simultaneous assessment of Caspase 3 activity and phosphatidylserine (PS) translocation

Spermptotic changes were detected in spermatozoa with the use of Annexin V 674 conjugate (Molecular Probes, Leiden Holland), which detects the translocation of phosphatidylserine (PS) from the inner to the outer leaflet of the plasma membrane associated with membrane changes related to sperm processing. Both stains were combined in a multiparametric test and evaluated by FC[8]. A final concentration of 5 × 10 ^6^ spermatozoa/ml was obtained by adding 10μL of diluted spermatozoa to 990 μL of Anexin Binding Buffer. Then samples were loaded with Hoechst 334242 (0.3μM) and CellEVENT (2 μM) and incubated at r.t. for 15 minutes. Then samples were washed by a short centrifugation spin for 12” and suspended in 200 μl of Annexin binding-buffer (solution in 10 mM HEPES, 140 mM NaCl, 2.5 mM CaCl_2_, pH 7.4). To 200 μL of sample per essay, 5μL of Annexin V were added. After 15 minutes of incubation in the dark at room temperature, 400 μL of 1 × Annexin binding-buffer were added before reading in the flow cytometer.

### Detection of phosphorylated AKT (Ser^473^) in stallion spermatozoa

Stallion spermatozoa were washed in PBS and fixed in 2% paraformaldehyde in PBS at r.t. for 15 minutes; after fixation cells were washed twice with PBS and once with PBS 1% BSA, permeabilized for 30 min in 0.1% saponin in PBS-1% BSA. Then samples were stained with 2 μL/ml of phospho-AKT Alexa fluor 488 conjugate (cat n° 4071, Cell Signalling Technology) and incubated in PBS-1% BSA for 30 min in the dark at r.t. Samples were then washed in PBS and analyzed in the flow cytometer [15].

### Flow cytometry

Flow cytometry analyses were conducted using a Cytoflex^®^ flow cytometer (Beckman Coulter) equipped with violet, blue and red lasers and a MACSQuant^®^ VYB (Miltenyi Biotech) flow cytometer equipped with Yellow, violet, and blue lasers (561, 405, 488nm). The quadrants or regions used to quantify the frequency of each sperm subpopulation depended on the particular assay. Forward and sideways light scatter were recorded for a total of 50,000 events per sample. Gating the sperm population after Hoechst 33342 staining eliminated non-sperm events. The instruments were calibrated daily using specific calibration beads provided by the manufacturers. A compensation overlap was performed before each experiment. Files were exported as FCS files and analyzed using FlowjoV 10.4.1 Software (Ashland, OR, USA). Unstained, single-stained, and Fluorescence Minus One (FMO) controls were used to determine compensations and positive and negative events, as well as to set regions of interest as described in previous publications from our laboratory [15, 28, 29]

### Computational flow cytometry (t-SNE analysis)

Flow cytometry data are usually analyzed using a series of 2D plots and manual gating, however the increase in the number of parameters measured increase the number of 2D plots to display for every marker combination; for example a combination of four colors will require 30 2D plots. To overcome these problems computational methods to automatically identify populations in multi-dimensional flow cytometry data have been developed [30]. Using Flowjo v 10.5.3 compensated data of each multi-parametric assay described in material and methods were exported as FCS files from the flow cytometer, and loaded in Flowjo for computational analysis, data were downsampled, concatenated and single cell events analyzed. Flow cytometry data were analyzed using non linear dimensionally reduction techniques (t-SNE). This technique identifies clusters within multidimensional data without loosing single cell resolution [31, 32], allowing automatic gating of cells. Within the t-SNE maps generated heat maps were applied to identify the expression of specific markers

### Statistical analysis

Frozen semen samples were obtained from 6 different stallions. All experiments were repeated at least three times with independent samples (three separate ejaculates from each of the donor stallions). The normality of the data was assessed using the Kolmogorov-Smirnoff test. Paired t tests and One-way ANOVA followed by Dunnett’s multiple comparisons test was performed using GraphPad Prism version 7.00 for Mac, GraphPad Software, La Jolla California USA, www.graphpad.com. Overton cumulative histogram subtraction was also performed [33], to determine positivity in selected cytometry analysis; in brief this method determines that percent of the events that are considered to have positive florescence for the selected parameter by subtracting out the florescence of the control. Differences were considered significant when P < 0. 05, and are indicated as; * p<0.05 and ** p<0.01. Results are displayed as means ± SEM.

## RESULTS

### Rosiglitazone in the thawing media improves sperm motility

When the thawing medium was supplemented with rosiglitazone (75 μM), significant improvements in motility were seen after two hours of incubation (p<0.01). Both the percentages of total motile spermatozoa and linear motile spermatozoa showed significant improvements with the treatment (Fig 1 A and B.). Other concentrations of rosiglitazone tested had no effect.

**Figure 1.**
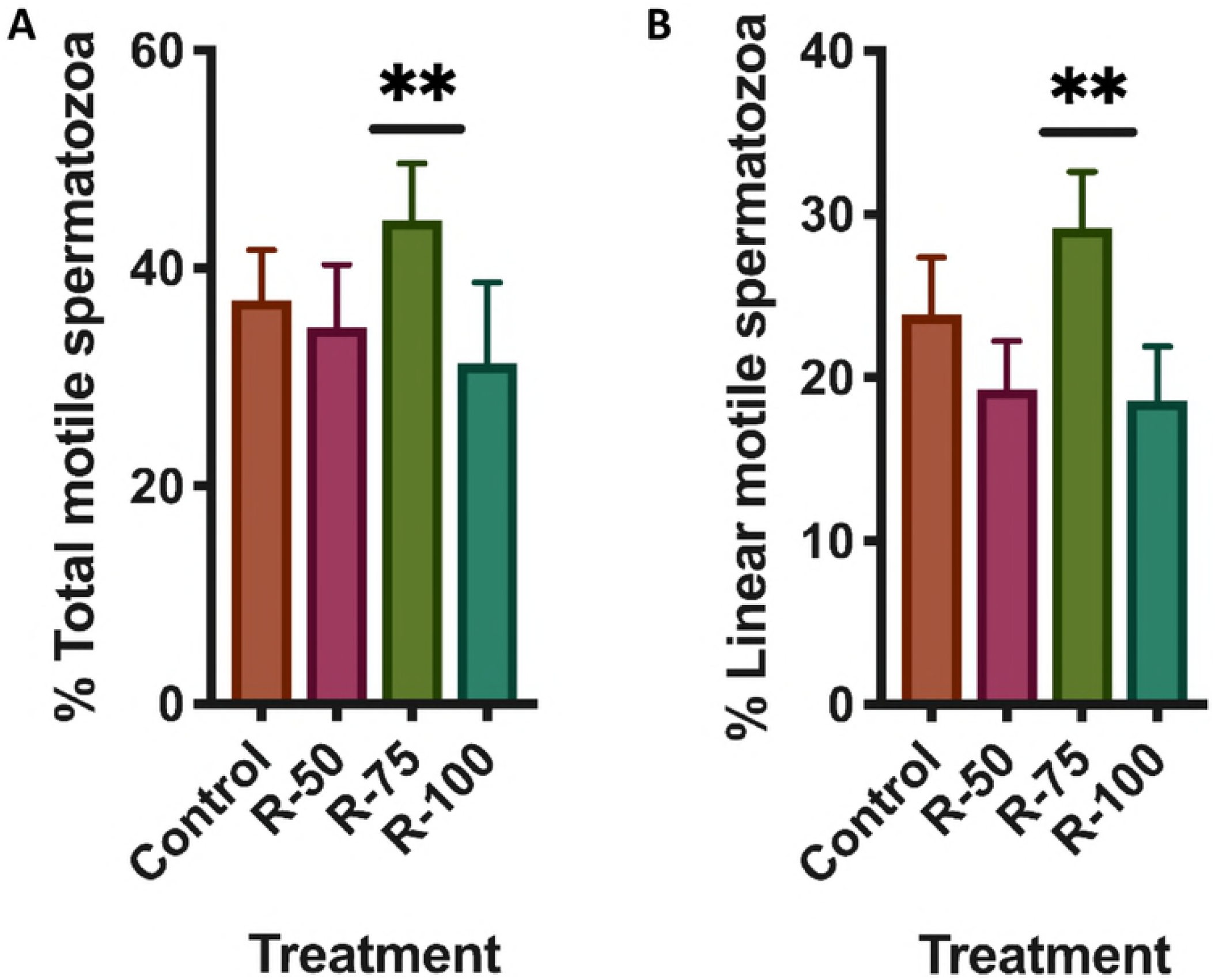
Effect of rosiglitazone added to the thawing media on stallion sperm motility. Samples were processed as described in material and methods, and supplemented in the thawing media with rosiglitazone (0, 50, 75 and 100μM) incubated at 37°C for two hours and motility evaluated using a computer assisted system (CASA). Rosiglitazone at 75 μM increased the percentage of total (A) al linearly motile (B) spermatozoa (*P*<0.01)

### Rosiglitazone enhances mitochondrial function in thawed stallion spermatozoa

Mitochondrial impartment is a hallmark of thawed stallion spermatozoa [8, 19, 34, 35], and also is considered an early event in spermptosis [8]. In order to determine if rosiglitazone is able to improve mitochondrial function, thawed stallion spermatozoa were incubated in presence of rosiglitazone (75 μM) and after 1 and two hours of incubation the mitochondrial function of surviving spermatozoa was investigated using JC-1. Since the production of the superoxide anion (0_2_•^-^) is a byproduct of oxidative phosphorylation in the mitochondria [36, 37], at the same time the production of 0_2_•^-^ was simultaneously investigated. Rosiglitazone significantly increased the mitochondrial potential of surviving spermatozoa at every moment of incubation considered (Fig 2 A-D). Increased mitochondrial activity estimated as increased presence of JC-1 aggregates [38] occurred without concomitant increases in the production of 0_2_•^-^ after 1 hour of incubation (Fig.2 E), but there was a significant increase in 0_2_•^-^ in supplemented samples after 2 hours of incubation at 38°C (Fig. 2 F). When the analysis was performed on a cell by cell basis of the whole sperm population, the heat map generated after the t-SNE analysis showed evident changes indicating that rosiglitazone increased mitochondrial activity (estimated as the number of JC-1 aggregates) as compared with controls (Fig 2 a-b) in the whole sperm population, although in different degrees. Also to identify the major source of 0_2_•^-^, a heat map was generated for superimposing the APC channel over the JC-1/H33342 2D plot, showing that the major production of 0_2_•^-^ occurred in more active mitochondria (Fig, 2 c).

**Fig 2.** Effects of rosiglitazone added to the thawing media on mitochondrial function of stallion spermatozoa after thawing. Frozen stallion semen was thawed and processed as described in material and methods. Split samples were supplemented with rosiglitazone (0 and 75 μM) and mitochondrial functionality was investigated after 1 and 2 hours if incubation. A and C, percentage of spermatozoa showing orange fluorescence after JC-1 staining, B and D, mitochondrial functionality expressed as the mean fluorescence intensity in the PE channel indicative of JC-1 aggregates (high mitochondrial potential), E and F production of superoxide after 1 and 2 hours of incubation at 37°C. Results are presented as means ± SD. * *P*<0.05, ** *P*<0.01. In a and b the t-SNE map after computational analysis is seen, in the t-SNE map each point represent individual spermatozoa in the sample, the heap map applied to the t-SNE map shows increased PE fluorescence (jc-1 aggregates) in rosiglitazone treated samples. In c a representative 2D plot after JC-1/H33342 is presented, A live spermatozoa with high ΔΨm, B live spermatozoa, C, dead spermatozoa. In d a heat map overlay of the APC channel (production of superoxide) is shown over the 2D plot depicted in c, maximum production of superoxide is present in live sperm showing high ΔΨm (orange events in the plot).

### Rosiglitazone does not modify the oxidation reduction potential (sORP)

In order to discriminate if the increased production of superoxide is just caused by intense mitochondrial activity [36], or is a sign of oxidative stress; the oxidation-reduction status of the samples was investigated. No changes were observed in the static oxidation reduction potential (sORP), or in the total antioxidant capacity in supplemented samples (Fig 3).

**Fig 3.** Effect of rosiglitazone added to the thawing media, on sORP (mV/10^6^ sperm) (A) that is the integrated measure of the existing balance between oxidants and reductants and (B) antioxidant capacity reserve cORP (μC).

### Rosiglitazone reduces caspase 3 activation without changes in phosphatidylserine transposition

Since its have been reported that caspase activation triggers sperm senescence [15], we studied the effect of rosiglitazone on caspase 3 activation; when thawing media was supplemented with rosiglitazone a significant decrease of >25% respect initial values in controls after 1 and 2 hours on incubation, occurred (P<0.05, P<0.01) (Fig. 4A-B). The study of the t-SNE map also showed a decrease in caspase 3 activation (Fig 5 A-B). Phosphatidylserine transposition increased over the treatment period, however no changes induced by rosiglitazone were observed (data not shown).

**Fig 4.** Changes in caspase 3 after rosiglitazone supplementation of the thawing media. Commercial frozen doses of stallion sperm were thawed and processed as described in material and methods. Split samples were incubated in presence of rosiglitazone 0 and 75 μM and caspase 3 activity flow cytometrically determined. Data represent percent changes respect controls after 1 hour (A) and two hours (B) of incubation and are expressed as means ± SD, * P<0.05

**Fig 5.** Computational cytometry analysis (t-SNE) graphics. including heat maps are presented showing the effect of 75 μM rosiglitazone supplementation of caspase 3 activity in thawed stallion spermatozoa. A, control samples, each point represent an individual spermatozoa, as seen in the heat map, almost half of the population shows high caspase 3 expression (orange color). B Samples supplemented with 75 μM rosiglitazone, as seen in the heat map, caspase 3 expression is reduced, only a small population still present high caspase 3 (red circle).

### Rosiglitazone phosphorylates Akt

Previous findings from our laboratory linked the dephosphorylation of the Akt and the activation of caspase 3 in ejaculated stallion spermatozoa [15]. We hypothesized that rosiglitazone may be linked to Akt phosphorylation. Incubation of thawed stallion spermatozoa in presence of rosiglitazone maintained phosphorylated Akt after two hours of incubation at 37°C in comparison with untreated controls (Fig. 6B and 7).

**Fig 6.** Effect of rosiglitazone on Akt phosphorylation (***Ser** ^473^*) on stallion spermatozoa. Commercial frozen doses of stallion sperm were thawed and processed as described in material and methods. Split samples were incubated in presence of rosiglitazone 0 and 75 μM and Akt phosphorylation measured after 1 (A) and 2 hours (B) of incubation at 38°C. Data represent percent changes respect controls and are expressed as means ± SD * P<0.05.

**Fig 7.** Representative overlay cytograms of the p-Akt assay after 1 hour (B) and 2 hours of incubation (D). To calculate expression of the different germ cell markers we used the population comparison analysis available in FlowJo, version 10.4.1 (TreeStar, OR, USA). This analysis uses the overton cumulative histogram subtraction algorithm (Overton, 1988), and overlaps histograms of control (isotype control) and sample, allowing subtraction of control to calculate the percentage of positive cells in the sample (percentage of cells showing increased expression respect controls)

### Inhibition of Akt, PPARγ and AMPK abolished the reduction of caspase 3 activation induced by rosiglitazone

The effects of rosiglitazone can be mediated by the PPARγ receptor and or by the phosphorylation of the AMPK. In both cases Akt can be phosphorylated [23]. To test this hypothesis samples were pre-incubated in presence of Akt1/2 inhibitor (30μM), GW9662, inhibitor of PPARγ (10μM) and dorsomorphin, inhibitor of AMPK (100μM) and then incubated in presence of rosiglitazone 75μM. As seen in the previous experiment rosiglitazone reduced caspase 3 activation, effect that was abolished in presence of the Akt1/2 inhibitor, GW9962 and dorsomorphin both after 1 and two hours of incubation (fig. 8 A-B).

**Fig 8.** Effects of AKT ½ Kinase inhibitor, dorsomorphin (AMPK inhibitor) and GW9662 (PPAR γ inhibitor) on caspase 3 inhibition after rosiglitazone treatment. Thawed semen doses were processed as described in M and M, and incubated in presence of rosiglitazone (0 and 75 μM) or rosiglitazone 75 μM plus Akt ½ kinase inhibitor 30μM, Rosiglitazone 75 μM plus GW9662 10μM or rosiglitazone 75μM plus dorsomorphin 100μM. After 1 and 2 hours of incubation caspase 3 activity was determined using flow cytometry. Results are presented as means ± SD. * P<0.05 A) changes after 1 hour of incubation, B) Changes after 2 hours of incubation

## DISCUSSION

In this study we aimed to determine whether the quality of frozen semen can be improved after thawing. Traditional approaches to improve sperm survival after freezing and thawing, focus on the improvement of extenders, sperm selection pre-freezing or post thawing, and freezing and thawing rates [5, 39–45]. Few studies have focused on the development of methods to improve sperm quality after the thawing phase; moreover few studies have addressed the biology of thawed spermatozoa. Our results show that the functionality of thawed stallion spermatozoa can be improved activating pro-survival pathways; particularly mitochondrial function can be significantly improved.

Recent studies point to stallion spermatozoa as highly dependent of intracellular thiols for their proper functionality [12, 46, 47], and also highly dependent of oxidative phosphorylation in the mitochondria as the main source of ATP for motility, but mainly for maintenance of membrane functionality [18, 35, 37, 48, 49]. These facts have important implications for the selection of more fertile spermatozoa [37] and sperm conservation [17, 48–50]. The thawed spermatozoa are characterized by compromised mitochondrial function [8, 14] and an unstable redox status, leading rapidly to oxidative stress [1, 14, 51]. We aimed to induce metabolic flexibility to improve the functionality of thawed spermatozoa, a strategy that has proven successful in the conservation in stallion spermatozoa in refrigeration [21]. Moreover we studied the potential mechanisms behind this improvement. The PPAR γ agonist rosiglitazone induced clear improvements in mitochondrial function, and reduced caspase 3 activity, these effects also linked to increased phosphorylation of Akt. Previous reports indicate the importance of Akt phosphorylation in sperm function [15, 16, 52, 53], and recently a link between PPARγ agonists and Akt phosphorylation in human [22] and pig spermatozoa [23] has been reported. Moreover strategies to maintain Akt phosphorylated in spermatozoa have proven to be successful in human sperm cryopreservation [52, 54]

Treatment described in our experiment was able to keep Akt phosphorylated in thawed spermatozoa. In addition experiments with Akt inhibitors further supported the link between the phosphorylation status of the Akt and sperm functionality as previously reported for human [16, 22] and equine spermatozoa. An important aspect of our experiment is that Akt can be maintained phosphorylated for longer periods in thawed sperm, suggesting that thawed sperm, although being exposed to extremely stressful osmotic and temperature conditions during the procedure, still retains ability to delay pro-death pathways. This is an interesting finding since opens new clues to develop strategies to improve the quality of frozen sperm after thawing. The prevalence of pro survival or pro-death pathways is probably linked to the redox status of the cell [13]. In this regard Akt can improve mitochondrial function in different cellular models, independently of transcriptional activity [55], supporting the proposing mechanism here described of enhanced mitochondrial function after PPARγ agonist treatment in spermatozoa. Further supporting this hypothesis, treatment of stallion spermatozoa with Akt inhibitors prevented improvements after rosiglitazone treatment, as did inhibition of PPARγ and AMPK. However inhibitor treatments indicated that in stallions most of the effect may be related through AMPK activation, firstly this effect has been previously reported in stallion semen maintained in the liquid state [21], and secondly, the PPARγ inhibitor was less efficient reverting rosiglitazone effects. More interestingly rosiglitazone enhanced mitochondrial function while maintained redox homeostasis; although increased superoxide production was observed after two hours, the oxidation reduction potential sORP did not change, suggesting that, as previously reported [56], increased production of superoxide may be an indicator of intense mitochondrial activity.

Cryopreservation causes a dramatic depletion of intracellular thiols in spermatozoa leading to unstable redox status rapidly leading to oxidative stress and cellular senescence [57], this form of sperm demise is characterized by activation of caspase 3. Interestingly, this activation, as described in our experiment can be delayed, thus increasing the lifespan of the spermatozoa. Beneficial effects of rosiglitazone are linked to the induction of metabolic flexibility in spermatozoa, using more efficiently glycolysis [21] and β oxidation of fatty acids [58] as sources of energy. Also as revealed in our experiment this pathway improves the efficiency of mitochondrial function. Mitochondrial function is considered a hallmark of functional spermatozoa [59–61], and more fertile stallion samples show more active mitochondria [37]; these reports support the concept that the quality and probably fertilizing ability of thawed samples can be modulated after thawing.

In conclusion, thawed stallion spermatozoa can be improved post thaw through mechanisms that maintain Akt phosphorylated, a process that may involve AMPK and PPARγ activation. Moreover these findings may have practical application to improve the quality of thawed samples independently of the initial freezing protocol.

## Acknowledgements

The authors received financial support for this study from the Ministerio de Economía y Competitividad-FEDER, Madrid, Spain, grants AGL2017-83149-R, Junta de Extremadura-FEDER (IB16030 and GR18008).

